# The membrane curvature inducing REEP1 proteins generate a novel ER-derived vesicular compartment

**DOI:** 10.1101/2023.12.19.572386

**Authors:** Yoko Shibata, Emily E. Mazur, Buyan Pan, Sebastien V. Hernandez, Jiuchun Zhang, Tom A. Rapoport

## Abstract

The endoplasmic reticulum (ER) is shaped by abundant, membrane curvature-generating proteins that include the REEP family member REEP5. The REEP1 subfamily, consisting of REEP1-4 in mammals, differs in abundance and topology from REEP5. Mutations in REEP1 and REEP2 cause Hereditary Spastic Paraplegia, but REEP1-4’s function remains enigmatic. Here we show that the REEP1 proteins reside in a novel vesicular compartment and identify features that determine their localization. Mutations in REEP1 proteins that compromise curvature-inducing activity, including those that cause disease, relocalize the proteins to the bulk ER. These mutants interact with wildtype proteins to retain them in the ER, consistent with their autosomal-dominant disease inheritance. REEP1vesicles contain the fusogen atlastin-1, but not general ER proteins. We propose that REEP1 proteins generate these vesicles themselves by budding directly from the ER, and that they cycle back to the ER by atlastin-mediated fusion. The vesicles may serve to regulate ER tubule dynamics.

## Introduction

The endoplasmic reticulum (ER) is an essential organelle in all cells, consisting of a dynamic, continuous membrane network of narrow tubules and planar sheets^1–5^. The relative abundance of tubules and sheets varies among cell types and even within a single cell, as exemplified by neurons, where dendrites and cell bodies contain both sheets and tubules, but axons are filled with only narrow tubules^6^. The high membrane curvature of tubules and sheet edges is stabilized by two distinct, conserved protein families, the reticulons and a branch of the REEPs (receptor expression-enhancing proteins) that in mammals includes REEP5^7–9^. The reticulons and REEP5 have redundant functions in shaping the ER. They are abundant in all eukaryotic cells and share a similar structure, consisting of four transmembrane segments (TMs), followed by an amphipathic helix near the C-terminus (APH-C). Both features are required to generate high membrane curvature^10–13^. The shaping and dynamics of the ER also requires a conserved fusion GTPase, called atlastin (ATL) in mammals. These membrane-bound GTPases mediate the tethering and fusion of ER tubules to form the three-way junctions of the polygonal network^14–19^.

The REEP family contains another branch, which in mammals consist of REEP1-4 (called REEP1 proteins). REEP1 proteins are highly conserved^20,21^, but their cellular function remains enigmatic. They have been implicated in ER morphology^20^, but they likely play a limited role in tubule formation, as they are much less abundant than the reticulons and REEP5. They have also been linked to lipid droplet formation^22,23^, mitotic ER dynamics^24^, nuclear pore formation^25^, and most recently, to macroautophagy in fission yeast^21^. REEP1-4 share high sequence similarity amongst each other and differ from the REEP5 subfamily in structure, as they are predicted to lack the N-terminal region, including the first TM, and to contain a disordered C-terminal tail^21^. The remaining three TMs are predicted to have a similar structure as the last three TMs of REEP5 and to be followed by an APH-C (**Fig. 1a**). Like REEP5, REEP1 proteins homodimerize through the TM domain, and both the TMs and APH-C are required for membrane curvature generation^21^.

**Figure 1.**
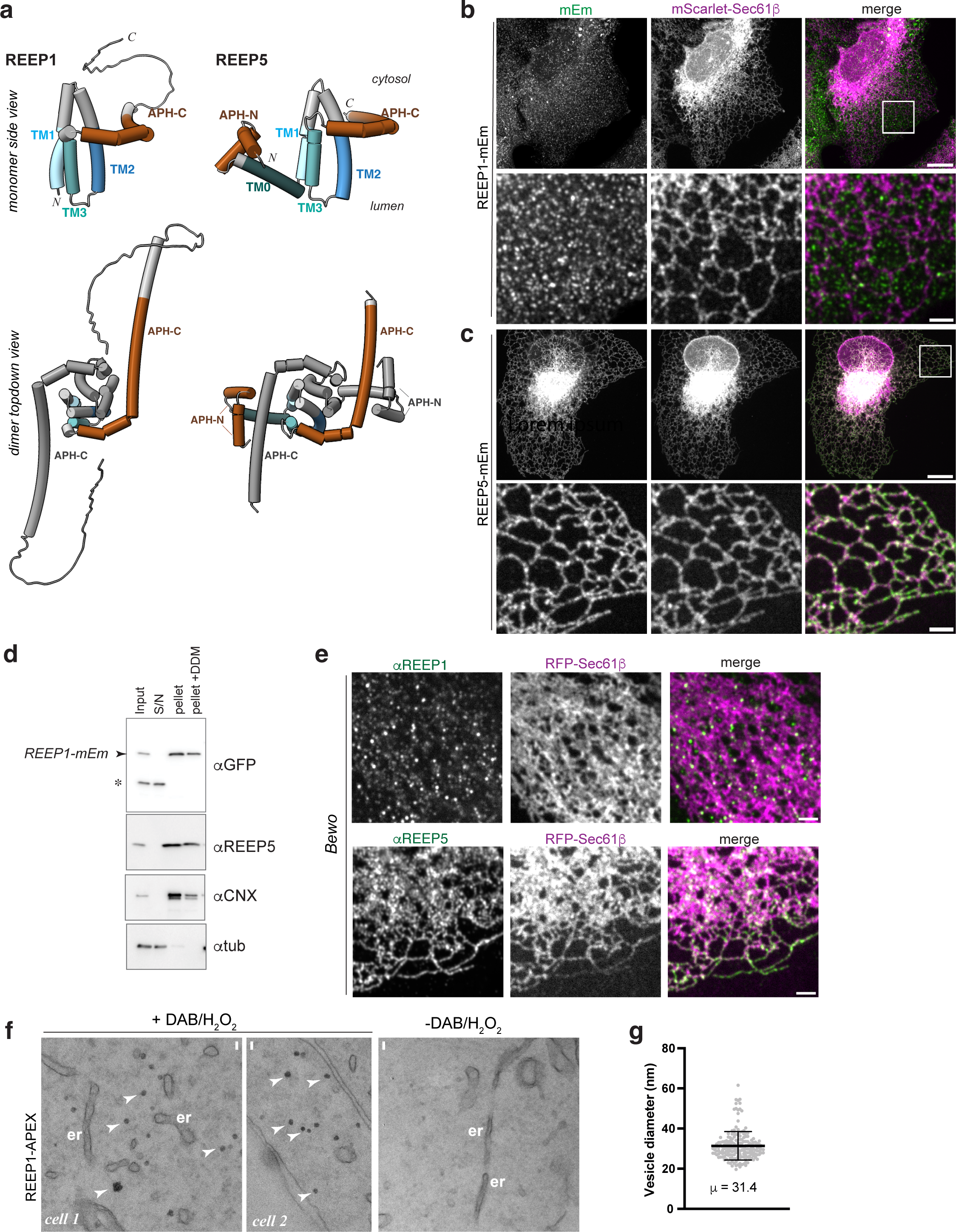
REEP1 localizes to vesicles in mammalian cells. **a**, Comparison of human REEP1 and REEP5 proteins. Shown are Alphafold-predicted models of the monomer and homodimer of each protein. Transmembrane (TM) segments are numbered. TMs are colored in blue and amphipathic helices at the N- or C-termini (APH-N or -C, respectively) in sienna. The second monomer in the dimer is shown in grey. **b**, U2OS cells stably co-expressing REEP1-mEmerald (REEP1-mEm) and the general ER marker mScarlet-Sec61β were imaged using confocal fluorescence microscopy. Bottom panels show magnifications of the boxed region. Scale bars of whole cells, 10 µm; magnifications, 2 µm. **c**, As in b, but with cells stably expressing REEP5-mEmerald (REEP5-mEm) and transiently transfected with mScarlet-Sec61β. **d**, Lysates from U2OS cells stably expressing REEP1-mEm were fractionated by ultracentrifugation into a supernatant (S/N) and a membrane pellet. A membrane extract was generated by treatment of the pellet with 1% DDM detergent followed by centrifugation. Samples were analyzed by SDS-PAGE and immunoblotting with antibodies against mEmerald (αGFP), endogenous REEP5 (αREEP5), the integral membrane protein calnexin (αCNX), and the soluble marker alpha tubulin (αtub). Input and S/N are 25% of the membrane fractions. **e**, Endogenous REEP1 (αREEP1, top row) and REEP5 (αREEP5, bottom row) localizations were analyzed by indirect immunofluorescence and confocal fluorescence microscopy in BeWo cells transfected with RFP-Sec61β. Scale bars, 2 µm. **f**, U2OS cells stably expressing REEP1-APEX were fixed, treated with diaminobenzidine and hydrogen peroxide (+DAB/H_2_O_2_), and analyzed by thin section electron microscopy. A sample without DAB/H_2_O_2_ treatment (-DAB/H_2_O_2_) is shown as a control. Arrowheads point to electron-dense, circular structures in +DAB/H_2_O_2_ samples. er, endoplasmic reticulum. Scale bars, 50 nm. **g**, Quantification of the diameters of REEP1-APEX structures shown in f. n, 186.

The REEP1 proteins are differentially expressed in distinct cell types. REEP1 and REEP2 are enriched in neurons, and mutations in these proteins are linked to hereditary spastic paraplegia (HSP) and distal hereditary motor neuropathy type Vb (HMN5B), two related, inherited neuropathies caused by axonal shortening of motor neurons^26–31^. HSP is also caused by mutations in ATL1^32^, consistent with the observation that ATLs interact with REEPs^14,20^. Given the physiological significance of this branch of the REEP family, it is important to identify the function of these proteins.

Here, we show that REEP1 proteins reside at steady state in a novel vesicular compartment and not the bulk ER, as hitherto assumed. The distinct localization is caused by the lack of the N-terminal domain found in the REEP5 subfamily. The vesicles appear derived from the ER and are likely generated by the REEP1 proteins themselves through their membrane curvature-inducing activity, rather than by conventional vesicle budding in the secretory pathway. ATL1 forms a physical complex with REEP1 and is found in these vesicles, allowing vesicles to recycle back to the ER. Disease-linked REEP1 mutations in the APH-C, which compromise curvature generation, lead to the retention of these proteins in the bulk ER. These mutants act dominantly by interacting with wildtype REEP1 proteins to retain them in the ER, thus inhibiting vesicle formation. Our results lead to a model in which REEP1 vesicles regulate the dynamics of ER tubules.

## Results

### Human REEP1 proteins localize to vesicles rather than the ER

We first compared the cellular localization of human REEP1 and REEP5 by stably expressing mEmerald-tagged versions (REEP1- or REEP5-mEm) in human U2OS cells. The bulk ER in these cells was labeled by expressing a mScarlet fusion of Sec61β (mScarlet-Sec61β). REEP1-mEm was found in punctae and short tubules distinct from the ER (**Fig. 1b**). In contrast, REEP5-mEm completely localized to the tubular ER (**Fig. 1c**); as observed previously^7,10^, REEP5-mEm was depleted in ER sheets and the nuclear envelope, consistent with its preference for high membrane curvature regions. The REEP1 punctae correspond to a membrane compartment and are not formed by protein aggregation, as REEP1-mEm sedimented with membranes and could be solubilized from the membranes with a mild detergent (**Fig. 1d**).

A punctate pattern with limited ER overlap was also seen when hemagglutinin (HA)-tagged REEP1-4 were transiently expressed in U2OS cells (**Supplementary Fig. 1a**). While it was previously reported that overexpressed REEP1 predominantly localizes to bundled ER tubules^20^, we observed bundled tubules only in cells with the highest levels of expression; moreover, the tubules did not contain the ER marker RFP-Sec61β (**Supplementary Figs. 1b, c**). The punctate localization is not caused by the tag, as the same pattern was also observed when untagged REEP1 was overexpressed in U2OS cells (**Supplementary Fig. 1d**). Quantification showed that REEP1 maintained its ER-independent punctate localization over a wide range of expression levels (**Supplementary Fig. 1e**). Overexpression of untagged REEP2 and REEP4 in U2OS localized similarly (**Supplementary Figs. 1f, g**). The distinct punctate localization of REEP1 proteins was also observed in Hela cells (**Supplementary Fig. 1h**).

Endogenous REEP1 also localized to punctae distinct from bulk ER proteins in Bewo cells, a cell type that expresses REEP1 at high levels (**Fig. 1e**). A similar punctate pattern was observed with endogenous REEP2 in SKN-SH cells (**Supplementary Fig. 1i**) and with endogenous REEP4 in U2OS cells (note that the REEP4 staining is absent in REEP4 knockout cells, **Supplementary Fig. 1j**).

To further characterize the punctate structures, REEP1 was expressed as a fusion with ascorbate peroxidase 2 (REEP1-APEX) in U2OS cells, and the localization of the protein was examined by thin-section electron microscopy after treatment with diaminobenzidine and H_2_O_2_ to form an electron-dense precipitate^33^. REEP1-APEX mostly localized to vesicles that have an average diameter of approximately 30 nm (**Figs. 1f, g**). Taken together, these data show that REEP1 proteins localize to a vesicular compartment, rather than the ER, as previously assumed.

### Vesicular localization of REEP1 proteins depends on the membrane-curvature generating APH-C

We next tested whether the APH-C affects the cellular localization of REEP1, as this region is essential for the generation of high membrane curvature^21^ and many mutations linked to HSP map to it (see **Supplementary Table 1**). We found that the disease-causing APH-C deletion mutants (ι1113-201 and ι1102-139) did not localize to vesicles, but rather the bulk ER, where they overlapped with RFP-Sec61β (ι1113-201 and ι1102-139 vs. wt; **Fig. 2a, b** versus **Supplementary Fig. 1a**). Similarly, a disease-linked APH-C deletion mutant of REEP2 relocalized from vesicles to the ER (ι1111-252 vs. wt; **Fig. 2c** versus **Supplementary Fig. 1a**). The colocalization was confirmed by quantification of Pearson’s correlation coefficients (**Fig. 2d**).

**Figure 2.**
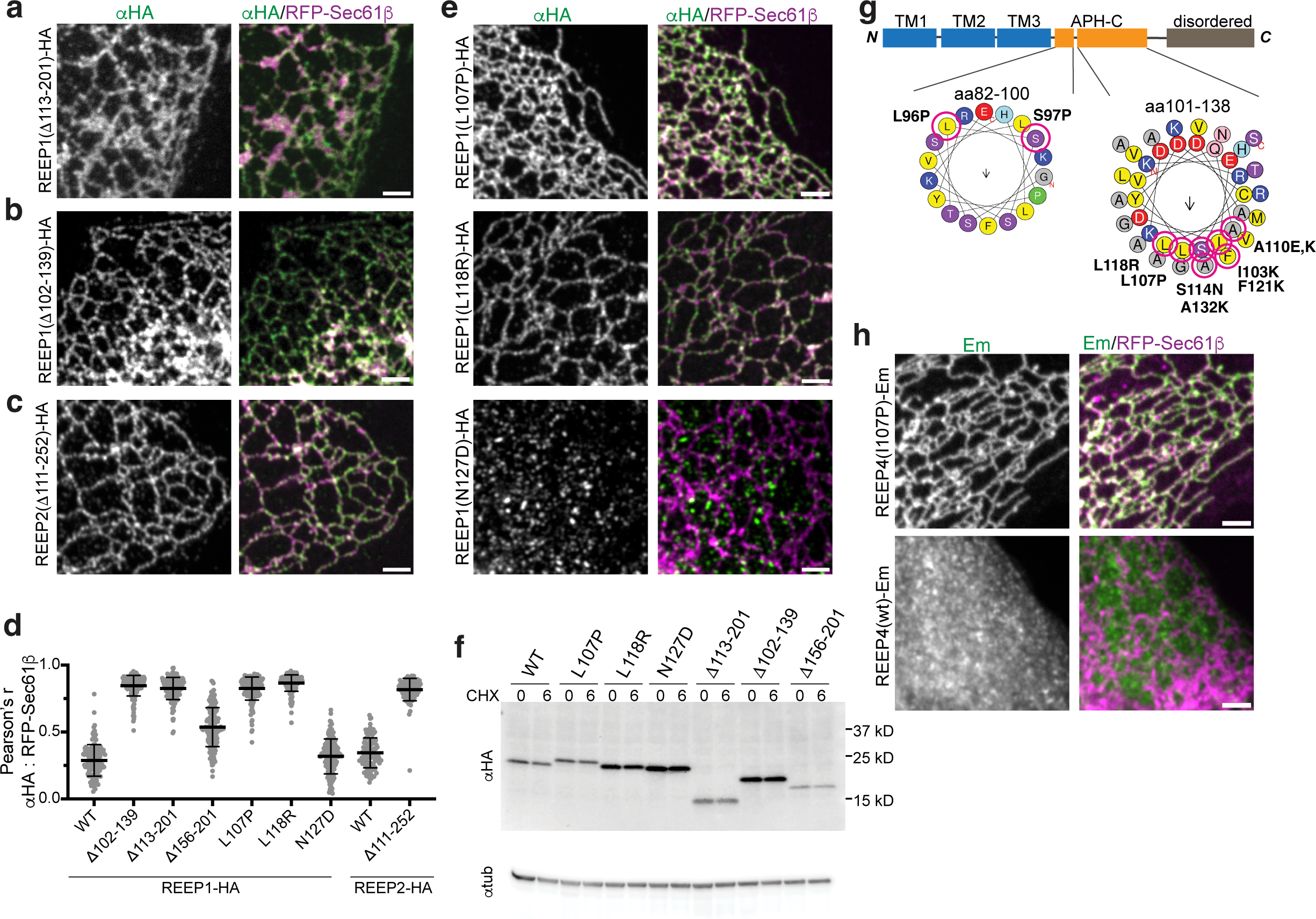
Vesicle localization of REEP1 proteins is dependent on their APH-C. **a,** A U2OS cell stably expressing RFP-Sec61β was transfected with HA-tagged REEP1 carrying the disease mutation Δ113-201, immunostained with anti-HA (αHA) antibodies, and imaged by confocal fluorescence microscopy. Scale bar, 2 µm. **b**, As in a, but in a cell transfected with the REEP1 disease mutant Δ102-139. **c**, As in a, but in a cell transfected with the REEP2 disease-linked mutant Δ111-252. **d**, Pearson’s correlation coefficients (r) measuring colocalization of RFP-Sec61β with wildtype or APH-C disease mutants of HA-tagged REEP1 or REEP2. Shown are means and standard deviations. n, 102-171 cells/sample. **e**, As in a, but in cells transfected with the REEP1-HA constructs carrying the L107P disease mutation, or putative disease mutations L118R or N127D (top, middle, and bottom rows, respectively). **f**, The relative stabilities of REEP1-HA APH-C mutants were analyzed by cycloheximide (CHX) chase. Cells transfected with wildtype or mutant HA-tagged REEP1 were collected at timepoint 0 or after CHX treatment for 6 h, and lysates were analyzed by immunoblotting with αHA antibodies. Anti-α tubulin serves as a loading control. **g**, Schematic of REEP1 domain organization and APH-C prediction. C-terminal REEP1 point mutations that cause relocalization to the bulk ER are circled in red. Amphipathic helices were predicted using heliquest (https://heliquest.ipmc.cnrs.fr/). The arrows indicate the hydrophobic moments. Note that all mutations are either helix breakers or reside along the APH-C’s hydrophobic side. **h**, REEP4 knockout U2OS cells transfected with RFP-Sec61β and stably expressing mEmerald (Em) fused REEP4(I107P) (top panels) or wildtype REEP4 (bottom panels) were imaged by confocal microscopy. Scale bars, 2 µm.

To analyze more systematically the features required in the APH-C for REEP1 vesicular localization, we introduced other deletions (**Supplementary Fig. 2a**) or mutated residues along the hydrophobic face of the APH-C into lysines (**Supplementary Fig. 2b**). Except for one mutation, A132K, all changes caused the relocalization of REEP1-HA to the bulk ER; the A132K mutant had an intermediate phenotype, displaying both punctate and ER localizations (**Supplementary Fig. 2b**). The deletion of the disordered C-terminal tail (ι1156-201) also caused the distribution to both punctae and ER (**Supplementary Fig. 2a**, quantification in **Fig. 2d**), suggesting that this region also plays a role in localization.

We next screened through disease-linked point mutations in the APH-C (see **Supplementary Table 1** for provenance). Several of these mutants localized to the bulk ER (L107P, L118R, **Fig. 2e**, quantification, **2d**; L96P, S96P, A110E, S114N, **Supplementary Fig. 2c**). Point mutations introduced into the hydrophilic side of the helix or into the disordered tail had no effect (N127D, **Fig. 2e**, quantification, **2d**; R124Q, G125S, T131A, T131I, V134M, G142R, R147I, P173L, K181T, **Supplementary Fig. 2d**). Cycloheximide-chase experiments showed that APH mutations causing relocalization of REEP1-HA to the ER do not overtly destabilize the proteins (**Fig. 2f**), making it unlikely that they are retained in the ER due to their misfolding. Remarkably, all point mutations that caused relocalization of REEP1-HA from vesicles to the ER are helix breakers or introduce positive charges into the hydrophobic face of the APH-C (depicted in **Fig. 2g**). A similar ER relocalization was also seen when an APH mutation was introduced into REEP4 fused to mEmerald (I107P, analogous to L107P of REEP1, **Fig. 2h**).

Taken together, our results show that the APH-C of REEP1 proteins is required for vesicular localization and that this localization is physiologically important. It is likely that the hydrophobic side of the APH-C dips into the cytosolic leaflet of the lipid bilayer, displacing phospholipid molecules and thereby generating high membrane curvature. The mutations would compromise lipid insertion and the generation of high membrane curvature. The relocalization of the REEP1 mutants also suggests that the REEP1-containing vesicles originate from the ER.

### The TM domain is important for REEP1 vesicle localization

Disease-causing mutations are also found in the membrane-spanning region of REEP1, where they often map to the highly conserved cytosolic loop between TM1 and TM2^26–28,31^ (**Supplementary Fig. 3a**). As reported previously^34^, many of these mutations cause the relocalization of REEP1-HA to lipid droplets (**Supplementary Fig. 3b-d**), although several also lead to redistribution to the bulk ER (P19L, S23F, T55K; **Supplementary Fig. 3c, d**). One mutation, D56N, had no effect on localization. All TM mutations that cause REEP1 relocalization to the bulk ER/lipid droplets were unstable, as shown by cycloheximide-chase experiments (**Supplementary Fig. 3e**). Thus, as suggested previously^28^, these mutations likely cause disease through loss of function. All mutants, except the D56N mutant, did not colocalize with REEP1-mEm stably expressed in the same cells (**Supplementary Fig. 3f-m**; quantification in **Supplementary Fig. 3n**), suggesting that they do not form dimers with the wildtype protein. The mutated residues map to predicted interaction regions within and between the REEP1 monomers (**Supplementary Fig. 3o**) and are likely essential for intramolecular folding and homodimerization.

### Features that cause the distinct localization of REEP1 and REEP5

Next, we asked what structural features cause the distinct localization of REEP1 and REEP5 to vesicles and the tubular ER, respectively. REEP5 has a unique N-terminal domain with two predicted APHs and an additional TM segment (**Fig. 1a; Fig. 3a**). Deleting this domain (Δ2-51) relocalized REEP5 to punctae that did not overlap with the ER marker RFP-Sec61β (Δ2-51versus wt, **Fig. 3b, c**; quantification in **Fig. 3d**). Interestingly, these REEP5Δ2-51 punctae did not colocalize with those containing REEP1 (**Fig. 3e**), suggesting that the REEP5 mutant and REEP1 generate their own distinct vesicles. As in the case of REEP1, the vesicular localization of the REEP5 Δ2-51 mutant depended on the APH-C: a double mutant with an inactive APH-C (Δ2-51; V156P) was found in the ER (**Fig. 3f**, quantification in **Fig. 3d**). When the N-terminal domain of REEP5, including its two APHs and the first TM, was fused to full-length REEP1, the chimera localized to the ER (**Fig. 3g, d**). Taken together, these results indicate that the unique N-terminus of REEP5 is a major determinant for ER tubule localization, clarifying why REEP1 and REEP5 have distinct localizations in mammalian cells.

**Figure 3.**
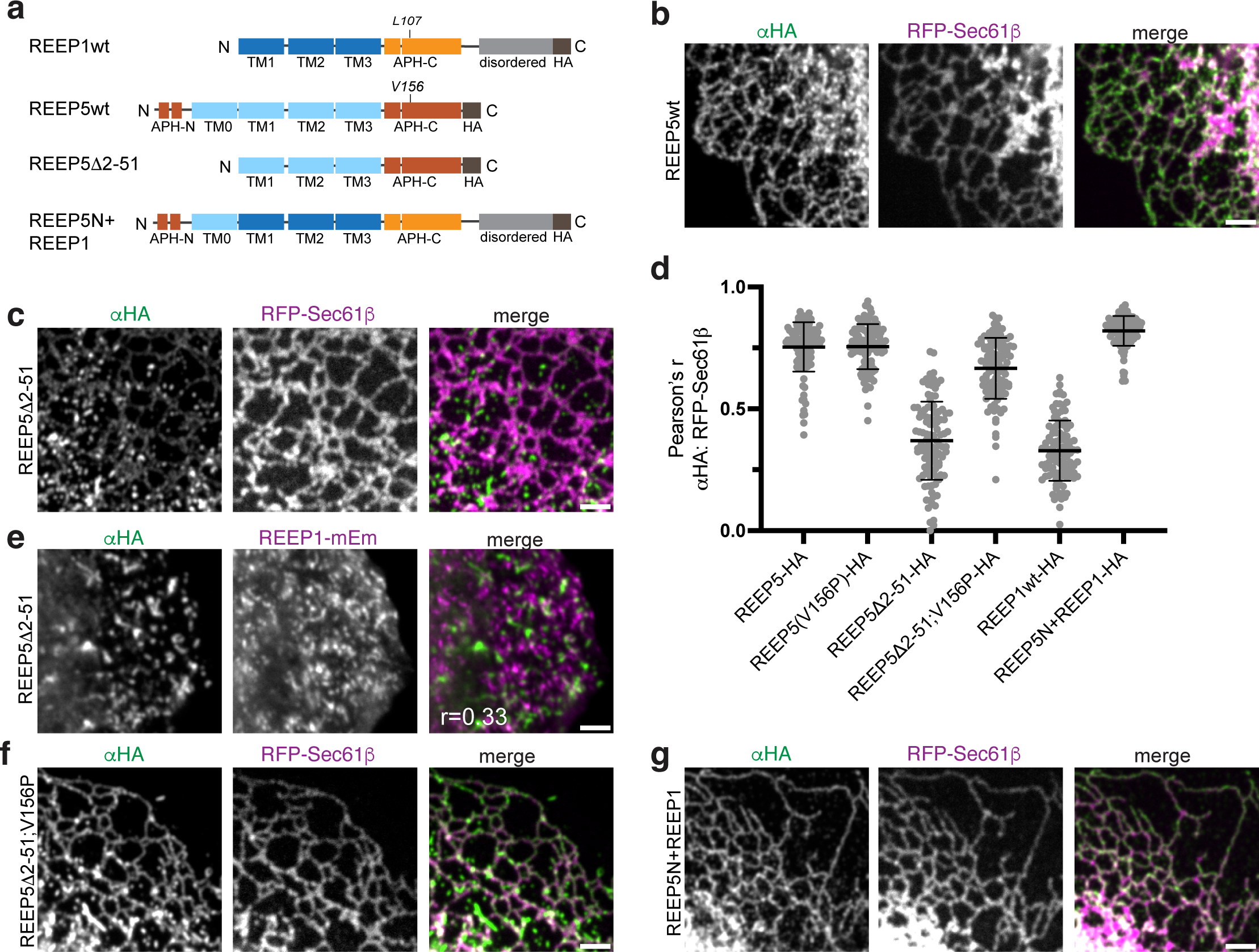
REEP5’s unique N-terminal domain is important for tubular ER localization. **a,** Schematic of REEP N-terminal deletion and chimera constructs used in b-g. Wildtype (wt) REEP1 is shown for comparison. Note that residue V156 in REEP5 corresponds to L107 in REEP1, which is mutated in disease. **b,** A U2OS cell stably expressing RFP-Sec61β was transfected with HA-tagged wildtype REEP5 (REEP5wt) and analyzed by immunostaining with αHA antibodies and confocal microscopy. Scale bar, 2 μm. **c**, As in b, but with transfection of a REEP5-HA construct lacking N-terminal residues 2-51 (REEP5Δ2-51). **d**, Pearson’s correlation coefficients (r) comparing localization of HA-tagged wildtype REEP1, REEP5, or REEP5 mutants with RFP-Sec61β in samples as analyzed in b, c, f, g. Colocalization coefficients were calculated between αHA and RFP. Shown are means and standard deviations. n, 101-111 cells/sample. **e**, As in c, but with a cell stably expressing REEP1-mEm. The Pearson’s correlation coefficient between αHA and mEm localization is shown in the inset. **f**, As in b, but with transfection of a REEP5Δ2-51 construct carrying an additional V156P mutation in the APH-C (REEP5Δ2-51; V156P). **g**, As in b, but with transfection of a chimera construct consisting of the first 51 residues of REEP5 fused to the N-terminus of wildtype REEP1 (REEP5N+REEP1).

### REEP1 mutants can drag wildtype REEP1-4 from vesicles into the ER

Given that REEP1 proteins form dimers through their TM domains, we wondered if the ER-localized REEP1 APH-C mutants could drag wildtype REEP1 proteins to the ER. We tested this possibility by co-expressing wildtype or mutant REEP1-HA with wildtype REEP1-mEm, using tandem constructs in which the two genes are separated by the ribosomal skip site P2A (**Fig. 4a**). As expected, wildtype REEP1-HA colocalized with REEP1-mEm in ER-independent punctae, and the TM domain mutant P19R did not change wildtype REEP-mEm localization (**Fig. 4b, c**). The ER-localized APH-C mutants of REEP1-HA (L107P or Δ102-139), however, redistributed REEP1wt-mEm to the bulk ER, although some ER-independent vesicles remained (**Fig. 4d, e**). Quantification of the colocalizations supports these conclusions (compare mEm: αKDEL Pearson’s values; **Fig. 4f**).

**Figure 4.**
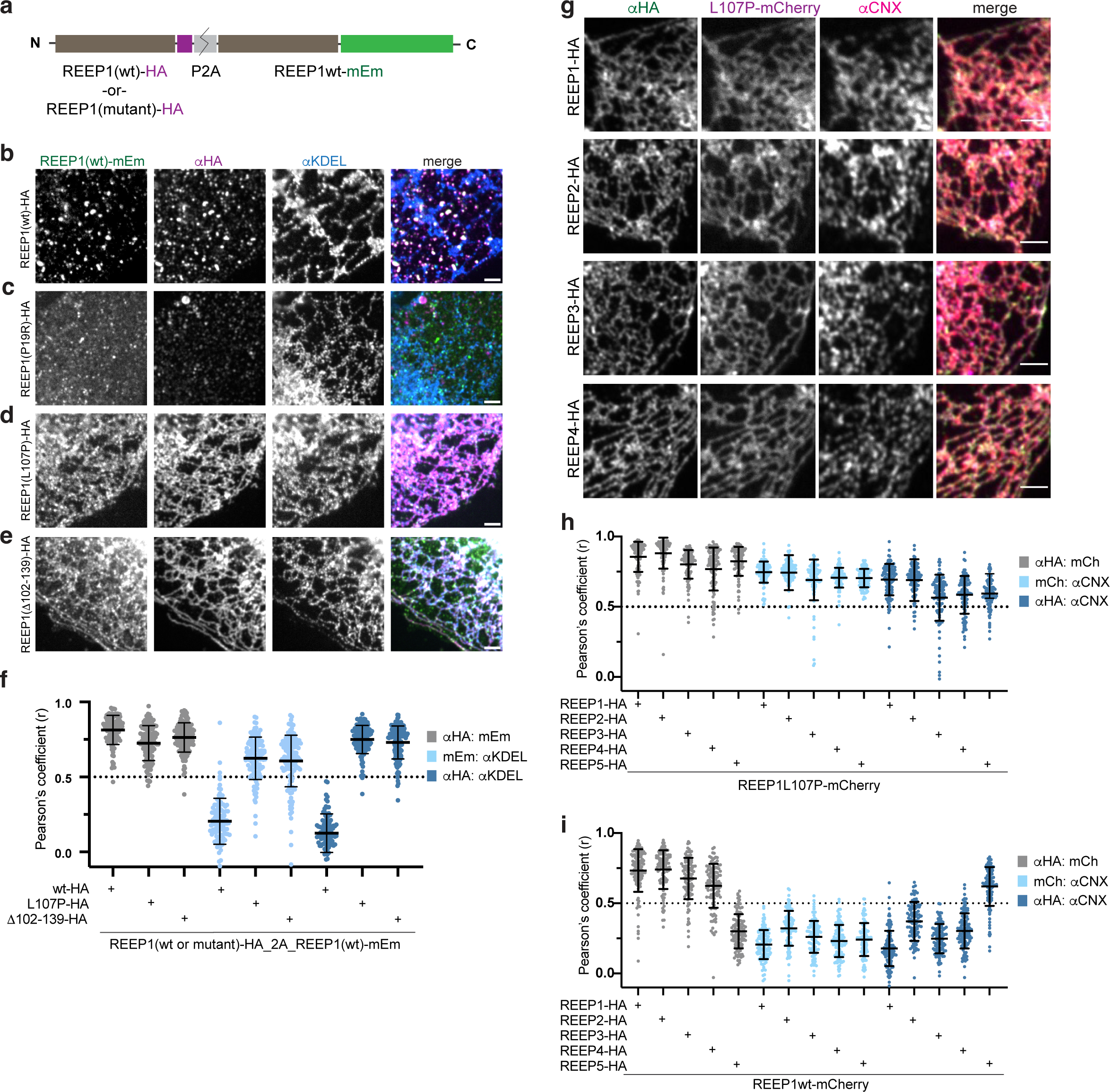
REEP1 APH-C mutants can ‘drag’ wildtype REEP1 proteins to the bulk ER. **a**, Schematic of tandem REEP1 constructs analyzed in b-f. Constructs encode for HA-tagged wildtype (wt) or mutant REEP1, the ribosomal skip site P2A, and wildtype REEP1 fused to mEmerald (mEm). **b**, A U2OS cell transfected with the REEP1wt-HA_P2A_REEP1-mEm construct was analyzed by immunostaining with αHA and αKDEL antibodies and imaged by confocal microscopy. Scale bar, 2 µm. **c**, As in b, but transfected with the REEP1(P19R)-HA_P2A_REEP1-mEm construct. **d**, As in b, but transfected with the REEP1(L107P)-HA_P2A_REEP1-mEm construct. **e**, As in b, but transfected with the REEP1(ι1102-139)-HA_P2A_REEP1-mEm construct. **f**, Pearson’s correlation coefficients (r) of U2OS cells expressing tandem REEP1 constructs, as in b, d, and e. The colocalizations of anti-HA to mEm, anti-HA to anti-KDEL, and mEm to anti-KDEL were quantified. Shown are means and standard deviations. n, 104-128 cells/sample. **g**, U2OS cells stably expressing REEP1(L107P)-mCherry were transfected with REEP1-HA, REEP2-HA, REEP3-HA, or REEP4-HA (top, second, third, or bottom row, respectively) and analyzed by immunostaining with αHA and αCNX antibodies and confocal microscopy. Scale bar, 2 µm. **h**, Pearson’s correlation coefficients of U2OS cells stably expressing REEP1(L107P)-mCherry and transfected with HA-tagged REEP1-5, as analyzed in g. Relative colocalizations between αHA and mCherry, mCherry and αCNX, and αHA and αCNX were quantified. n, 102-115 cells/sample. **i**, Pearson’s correlation coefficients as in h, but in cells stably expressing wildtype REEP1-mCherry. n, 101-115 cells/sample.

Given that the TM domain is highly conserved among REEP1-4, we next tested whether the disease mutant REEP1 L107P could also drag other REEP1 paralogs into the ER. To this end, we generated a stable cell line that expresses a mCherry fusion of REEP1 L107P (REEP1L107P-mCh). Like the HA-tagged mutant (**Fig. 2d**), the fusion protein localized to the bulk ER where it colocalized with endogenous calnexin (**Fig. 4g**). When wildtype REEP1-HA was transfected into these cells, it was dragged into the bulk ER, as were HA-tagged wildtype REEP2-4 (**Fig. 4g**; Pearson’s correlation quantification in **Fig 4h**). REEP5-HA localized to the tubular ER regardless of REEP1L107P-mCh (**Fig. 4h**). These data indicate that the REEP1 L107P mutant can interact with all members of the REEP1-4 subfamily. The mutant can retain the wildtype REEP1-4 proteins in the bulk ER, preventing their movement to the vesicular compartment and thereby acting in a dominant-negative manner. Our results offer an explanation for the autosomal dominant nature of APH-C disease mutations.

Heterodimerization of the REEP1-4 family members raises the possibility that they normally reside in the same vesicles. To test this idea, we transfected HA-tagged versions of wildtype REEP1-REEP4 into U2OS cells that stably express wildtype REEP1-mCh. All HA-tagged proteins colocalized with REEP1-mCh, but not with the ER marker calnexin (**Fig. 4i**). Thus, all REEP1 family members reside in the same vesicles. In contrast, REEP5-HA maintained its localization to the tubular ER and did not affect REEP1-mCh punctate localization (**Fig. 4i**), suggesting that REEP5 does not interact with the REEP1-4 subfamily.

### REEP1 vesicles comprise a novel compartment

We further characterized the REEP1 vesicles by testing colocalization with known cellular structures. The vesicles did not colocalize with F-actin (**Supplementary Fig. 4a**), but some aligned along microtubules (**Supplementary Fig. 4b**), although their distribution did not change with microtubule depolymerization (**Supplementary Fig. 4c**). We found no major colocalization of REEP1-mEm or -mCh with vesicles corresponding to early, late, or recycling endosomes (GFP-Rab5, -Rab7, or -Rab11, respectively), or to vesicles marked by the autophagosomal protein ATG9A^35^ (**Fig. 5a-d**; quantification in **Fig. 5e**). Additionally, no extensive colocalization was seen with lipid droplets (lipidtox), PIPERosomes^36^, phosphatidylinositol synthase (mCh-PIS)^36^, mitochondria (αTom20), or Golgi (αgiantin) (**Fig. 5f-h; Supplementary Fig. 4d, e**). There was also no significant overlap with ER exit sites (αSec31, **Fig. 5i**, quantification in **e**). REEP1 vesicle localization remained unaltered when ER-to-Golgi transport was inhibited with Brefeldin A^37^ or by incubation at 16 °C^38^ (**Supplementary Figs. 4f, g**). These results suggest that the REEP1 vesicles constitute a novel compartment.

**Figure 5.**
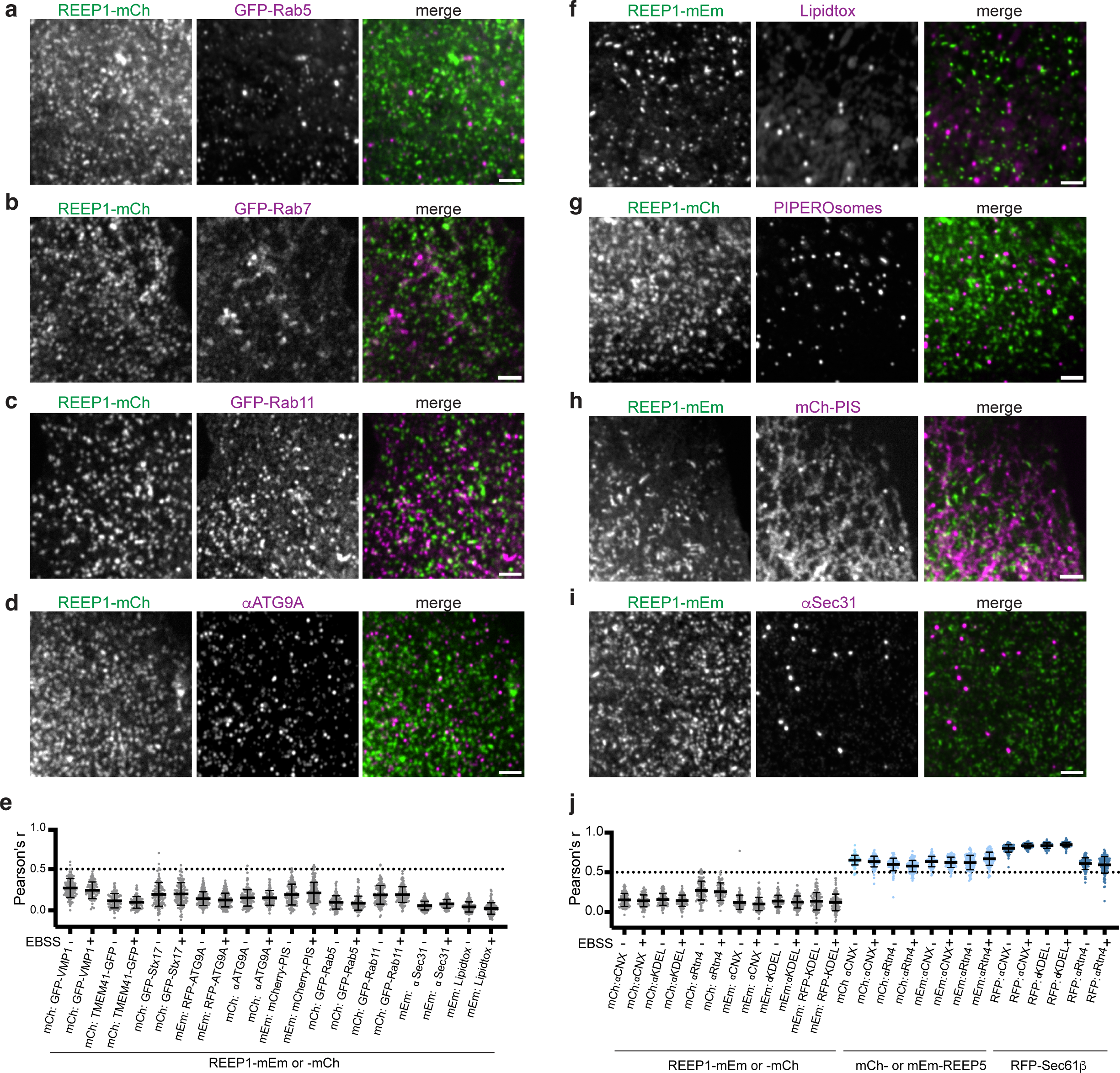
REEP1 vesicles do not colocalize with membrane-bound compartments. **a**, A U2OS cell stably expressing REEP1-mCherry and transfected with the early endosomal marker GFP-Rab5 was imaged by confocal microscopy. Scale bar, 2 µm. **b**, As in a, but with transfection of the late endosomal marker GFP-Rab7. **c**, As in a, but with transfection of the recycling endosomal marker GFP-Rab11. **d**, As in a, but analyzed by immunostaining with anti-ATG9A antibodies. **e**, Pearson’s analysis of REEP1-mCh or -mEm colocalization with various autophagy or vesicular markers in U2OS cells cultured in full or starved medium (-or + 30 min EBSS). See text for a description of markers. Note that REEP1 does not extensively colocalize with any marker tested. **f**, As in a, but in cells stably expressing REEP1-mEm and stained with the neutral lipid dye Lipidtox to visualize lipid droplets. **g**, As in a, but in cells coexpressing *Listeria* phospholipase C and the C1 tandem domains of protein kinase D fused to mEm to visualize PIPEROsomes. **h**, As in a, but in cells stably expressing REEP1-mEm transfected with mCh-PIS, which localizes to both the ER and ER-associated punctae. **i**, as in a, but in cells stably expressing REEP1-mEm and immunostained with anti-Sec31 antibodies. **j**, Pearson’s correlation coefficients comparing REEP1 localization to various ER markers. U2OS cells stably expressing REEP1-mCh or -Em, were analyzed as in e in either fed or starved conditions, and colocalized against the ER membrane protein calnexin (αCNX), an endogenous luminal marker (αKDEL), an overexpressed luminal marker ss-mScarlet-KDEL (RFP-KDEL), or an endogenous tubular ER marker (αRtn4). As controls, REEP5-mCh, REEP5-mEm, or RFP-Sec61β were also compared to αCNX, αKDEL, and αRtn4. Means and standard deviations are indicated. n, 74-152 cells/sample.

Given that the REEP1 ortholog Rop1 plays a role in macroautophagy in fission yeast, we further tested whether REEP1 changes its localization under starvation conditions in U2OS cells. The majority of REEP1-mEm and -mCh marked vesicles remained distinct from the ER even under starvation. They did not contain prominent endogenous ER membrane proteins (calnexin, Rtn4), endogenous luminal proteins (αKDEL), or overexpressed luminal RFP-KDEL (quantification in **Fig. 5h**). REEP1 vesicles also remained distinct from other membrane compartments or lipid droplets (**Fig. 5d**). A small fraction of stably expressed REEP1-mEm was found in starvation-induced phagophore membranes seen by the autophagy markers LC3 and DFCP1^39^ but lacking the lysosomal protein LAMP1 (**Supplementary Fig. 4h**). However, outside of the phagophore, REEP1-mEm vesicles did not colocalize with proteins involved in autophagy, including ATG9A, the lipid scramblases VMP1^40^ and TMEM41^41^, or the SNARE protein Stx17^42^ (**Fig. 5d**). These results indicate that a small fraction of REEP1 can be recruited to phagophores during starvation, but the majority of REEP1 proteins resides in unique vesicles.

### REEP1 localization is dependent on atlastin

Because curvature deficient REEP1 mutants localize to the ER, we wondered whether REEP1 would normally cycle between the ER and the novel vesicular compartment. Such cycling would require the fusion of the vesicles with the ER and implies that the REEP1 proteins are packed into the vesicles together with a fusogen. A likely candidate for the fusogen is the membrane-bound ATL GTPase, which typically mediates homotypic ER membrane fusion^14,15^ and has been reported previously to interact with REEP1 in co-immunoprecipitation experiments^20^.

We first tested whether REEP1 and ATL efficiently interact with one another, as this would facilitate co-packaging of the two proteins into budding vesicles. SBP-tagged REEP1 was expressed either alone or with FLAG-tagged ATL1 in 293 cells, the membranes were solubilized in the detergent digitonin, and the samples were subjected to immuno-isolation with FLAG antibodies followed by size-exclusion chromatography (**Fig. 6a; Supplementary Fig. 5**). REEP1 and ATL1 indeed formed a complex in which both proteins are present in approximately equal concentrations (**Fig. 6a**).

**Figure 6.**
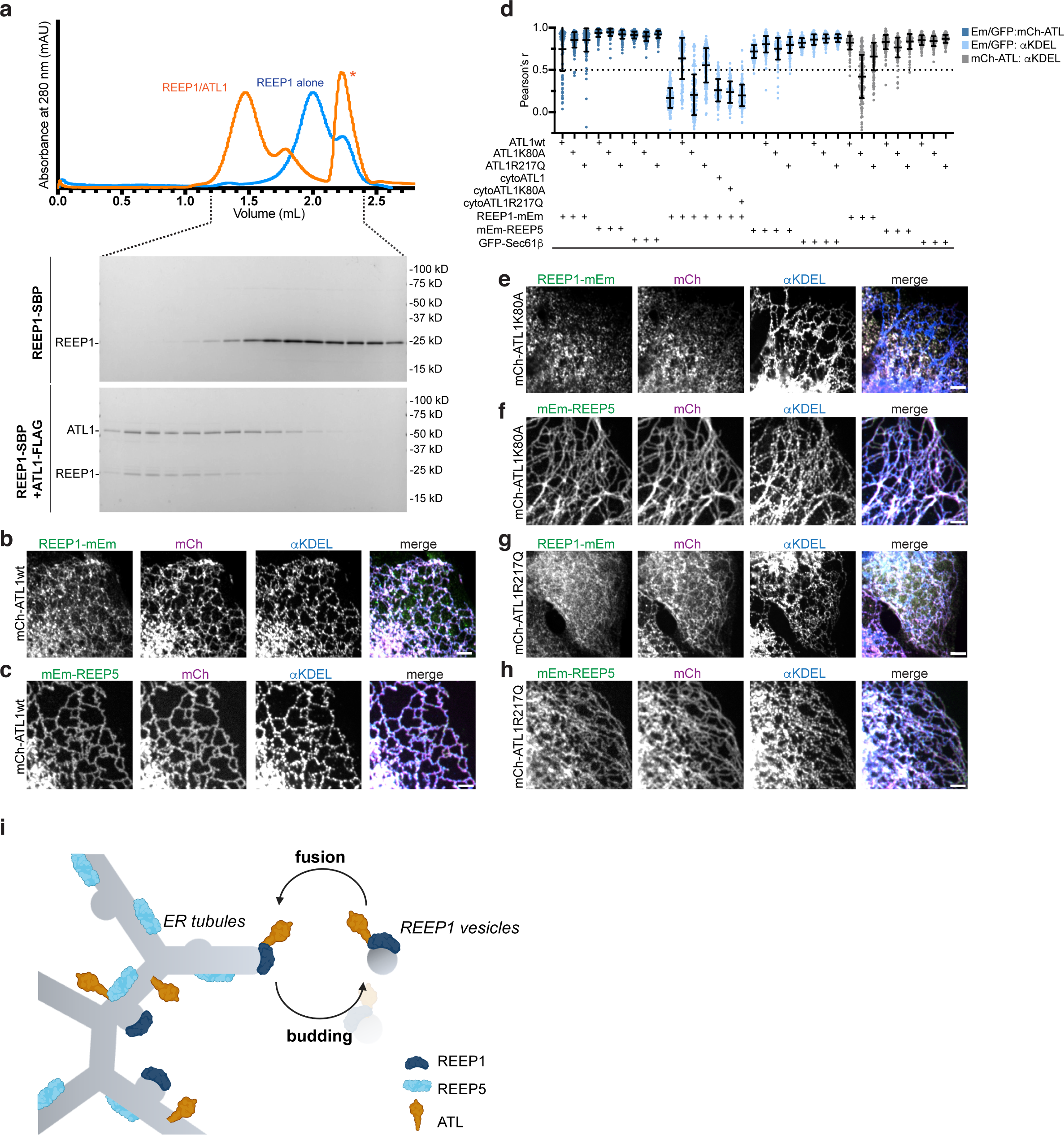
ATL1 GTPase colocalizes with REEP1. **a**, SBP-tagged REEP1 (REEP1-SBP) was expressed alone or with FLAG-tagged human atlastin-1 (ATL1-FLAG) in Expi293 cells and purified by solubilization in digitonin and binding to streptavidin-(REEP1-SBP) or anti-FLAG (ATL1-FLAG/REEP1-SBP) beads. The eluted material was analyzed by gel filtration. The peak marked with an asterisk corresponds to the 3xFLAG peptide used to elute ATL1-FLAG from anti-FLAG beads. Fractions from gel filtration elutions were analyzed by SDS-PAGE and Coomassie blue staining. **b,** A U2OS cell stably co-expressing REEP1-mEm and mCherry-fused wildtype ATL1 (mCh-ATL1wt) was immunostained with αKDEL antibodies and imaged using confocal microscopy. Scale bar, 2 µm. **c**, As in b, but in cells stably co-expressing mEm-REEP5 instead of REEP1-mEm. **d**, Pearson’s correlation analysis comparing the localizations of stably expressed mEmerald or GFP fusions REEP1-mEm, mEm-REEP5, or GFP-Sec61β to that of mCherry (mCh)-fused ATL1 wt or mutant constructs. Correlations between mEm (or GFP) and mCh, mEm/GFP and αKDEL, and mCh and αKDEL were quantified. Shown are means and standard deviations. n, 91-204 cells/sample. **e**, As in b, but in cells stably co-expressing the ATL1 GTP-hydrolysis mutant K80A (mCh-ATL1 K80A). **f**, As in c, but in cells stably co-expressing the ATL1 GTP-hydrolysis mutant K80A (mCh-ATL1 K80A). **g**, As in b, but in cells stably co-expressing the ATL1 nucleotide binding-deficient mutant R217Q (mCh-ATL1 R217Q). **h**, as in c, but in cells stably co-expressing the ATL1 nucleotide binding-deficient mutant R217Q (mCh-ATL1 R217Q). **i**, Model for REEP1 vesicle formation and recycling from the ER. See text for details.

To test whether ATL1 plays a role in REEP1 vesicle recycling, we generated stable U2OS cell lines co-expressing mCherry fused to wildtype ATL1 (mCh-ATL1wt) and mEmerald tagged REEP1, REEP5, or GFP-tagged Sec61β. Interestingly, when mCh-ATLwt and REEP1-mEm were co-expressed, they colocalized together, but REEP1-mEm became more ER-bound (**Fig. 6b**). mCh-ATL1wt maintained localization to the ER, as seen by co-staining with anti-KDEL antibodies, similar to when it was co-expressed with mEm-REEP5 or with GFP-Sec61β (**Fig. 6c, d**). Quantifications of Pearson’s coefficients confirmed these localization patterns (**Fig. 6d**). Thus, increasing levels of ATL moves REEP1 from vesicles to the ER.

Next, we performed similar experiments with an inactive ATL1 mutant which forms dimers that cannot undergo GTP hydrolysis^43^ (mCh-ATL1 K80A). When mCh-ATL1 K80A was co-expressed with REEP1-mEm, both proteins colocalized in ER-independent vesicles (**Fig. 6e**, quantification in **Fig. 6d**). In contrast, when mCh-ATL1 K80A was co-expressed with mEm-REEP5 or GFP-Sec61β, both proteins localized to the ER, as shown by co-staining with anti-KDEL antibodies (**Fig. 6f**, quantification in **Fig. 6d**). These results show that REEP1 can drag the inactive ATL1 K80A mutant from the ER into vesicles and suggest that the activity of ATL is required for the fusion of the vesicles with the ER.

An ATL1 mutant defective in dimerization (R217Q)^43^ behaved in an intermediate fashion, as it colocalized with REEP1-mEm in both the vesicles and the ER; its expression had no effect on the localization of mEm-REEP5 or GFP-Sec61β (**Figs. 6g, h**; quantification in **Fig. 6d**). REEP1 relocalization to the ER required the TM domain of ATL1, as the co-expression of ATL mutants lacking this domain (cytoATL1) left REEP1 in vesicles (**Fig. 6d**). Taken together, our results indicate that ATL1 and REEP1 are co-packaged into vesicles that bud from and fuse back to the bulk ER.

## Discussion

Here we show that a subfamily of the REEPs, REEP1-4, localizes to vesicles that do not contain markers of known compartments and therefore define a new organelle. We find it remarkable that after decades of cell biology, a novel compartment can still be discovered.

The vesicles likely form by the ability of REEP1 proteins to generate high membrane curvature, rather than by budding of COPII-coated vesicles from ER exit sites in the normal secretory pathway (see model in **Fig. 6i**). Indeed, all REEP1 mutations that compromise curvature generation localize the protein to the bulk ER and abrogate vesicle formation. The budding of REEP1-4 vesicles from the ER may be regulated by ATL, which also can generate high membrane curvature^44^. Co-packaging of these proteins into the same vesicles is facilitated by their ability to form near-stoichiometric complexes. Because the different members of the REEP1 subfamily can form hetero-dimers, all can co-segregate into the same budding vesicles.

Our experiments show that ATL1 is required for the fusion of REEP1 vesicles back to the ER; when ATL1 is inactivated by the K80A mutation, both the ATL1 mutant and REEP1 remain in vesicles. These results are consistent with the established fusion mechanism of ATL; ATL molecules in the ER-derived vesicles would tether with those in the ER and drive fusion between the two compartments via GTP hydrolysis. Cells must maintain a balance between the concentrations of REEP1-4 on the one hand and ATL on the other, as we found that moderate overexpression of wildtype ATL1 retains REEP1 in the ER. As ATLs are normally in excess to REEP1 proteins in tissue culture cells^45^, the abundance of these vesicles is likely lower than in our REEP1-overexpressing system.

The generation of this unique vesicular compartment is physiologically important, as mutations in REEP1 or REEP2 that cause neuronal diseases abolish the formation of the vesicles and relocalize the proteins to the ER. Many disease mutations map to the APH-C of these proteins, which likely reduce their ability to generate high membrane curvature. Because the mutants still form dimers, they can retain the wildtype proteins in the bulk ER, explaining their autosomal dominant nature. The fact that ATL1 is also commonly mutated in HSP is consistent with these proteins being involved in the same pathway. A genetic interaction between REEP1 and ATL1 is supported by experiments in mice, in which the deletion of REEP1 or the expression of the ATL1 K80A mutant alone had a relatively small effect, but the combination of both mutations caused strong neurological pathologies^46^.

In contrast to REEP5, most REEP1-4 molecules do not localize to the ER as hitherto assumed. Thus, despite both protein families being able to generate high membrane curvature, they have distinct localizations. Our analysis shows that the N-terminal region of REEP5, which includes two amphipathic helices and the first TM segment, is a major factor determining ER localization. Without this domain, REEP5 also localizes to vesicles, but these are distinct from REEP1 vesicles, supporting the idea that these proteins generate vesicles on their own. We speculate that the N-terminal domain allows REEP5 to generate anisotropic curvature that favors its localization to ER tubules, which have high curvature in cross-section but low curvature along the tubule length. REEP1 proteins, lacking this domain, would generate isotropic curvature that favors their localization to spherical vesicles, which have high curvature in all directions.

Assuming that REEP1 proteins induce isotropic curvature to form spherical vesicles, it seems likely that they bud from and can generate ER tubule tips, as these already have the shape of half-spheres (**Fig. 6i**). We propose that REEP1 proteins cooperate with ATL to generate dynamic tubule ends that can shrink and grow, adding an additional layer to how the ER tubular network remodels itself. In neurons, a deficiency of REEP1/2 or ATL1 would compromise the dynamics of ER tubules in axons of motor neurons and may cause the ER network to be more static or convert into nontubular structures^46^, leading to HSP and HMN5B neuropathies.

REEP1 proteins may also play a role independently of ER tubule dynamics, as our results indicate that a subpopulation of REEP1 is recruited to phagophores in starved tissue culture cells. In fission yeast, the REEP1 ortholog Rop1 is needed for the formation of phagophores, relying on its ability to generate high membrane curvature^21^. Whether Rop1 also cycles between the ER and vesicles in fission yeast remains to be tested.

In summary, our results lead to an extended model for how the tubular ER network is formed and remodeled (**Fig. 6i**). ER tubules themselves are generated by the reticulons and REEP5, which are some of the most abundant ER proteins and probably generate anisotropic high membrane curvature. REEP1 proteins would generate isotropic high membrane curvature and localize to ER tubule tips, where they cause vesicle budding and allow the tubules to shrink and grow. Finally, ATL is required for the fusion of ER membranes with each other and is involved in the budding and fusion of REEP1 vesicles.

## Acknowledgements

The authors thank the Nikon Imaging Center and the Electron Microscopy Facility, both at Harvard Medical School, for technical assistance and use of microscopes. We thank L. Clark for early experiments purifying REEP/ATL complexes; C. Blackstone, M. Davidson, N. Mizushima, R. Pagano, G. Voeltz, Do-Hyung Kim, E. Campeau, C. Vakoc, and H. Tukachinsky for plasmids; and M. Hoyer and D. Pellman for critical reading of the manuscript. S.V. H. and J. Z. were supported by the HMS Cell Biology Initiative for Genome Editing and Neurodegeneration. T.A.R. is a Howard Hughes Medical Institute Investigator. This article is subject to HHMI’s Open Access to Publications policy. HHMI laboratory heads have previously granted a non-exclusive CC BY 4.0 license to the public and a sublicensable license to HHMI in their research articles. Pursuant to those licenses, the author-accepted manuscript of this article can be made freely available under a CC BY 4.0 license immediately upon publication.

## Methods

### Plasmids and antibodies

All plasmids used and generated in this study are described in Supplementary Table 2. All gblocks were purchased from IDT and Gibson assembly reactions were performed with HiFi assembly mix (NEB). Site-directed mutagenesis was performed using Pfu Turbo polymerase (Agilent). All plasmids were sequence-verified using either Sanger sequencing of the inserted/mutated region or with nanopore whole plasmid sequencing.

All antibodies used in this study are listed in Supplementary Table 2.

### Mammalian tissue culture

All mammalian cell lines used and generated in this study are listed in Supplementary Table 2. All adherent cells were maintained at 37 °C and 5% CO_2_ in complete media supplemented with 10% FBS and penicillin-streptomycin (100 U/ml). Suspension Expi293 cells used for protein purifications were grown in Expi293 media (ThermoFisher) at 37 °C/8% CO_2_/80% humidity in an Infors Multitron shaker.

For stable cell line generation, U2OS cells were either transfected with the indicated plasmid or infected with lentiviral or retroviral particles. Positive cells were selected using antibiotics (0.5 μg/ml G418, 0.01 μg/ml puromycin, or 100 μg/ml blasticidin) to generate pooled lines. Clonal lines were isolated from pooled lines by plating single cells into 96-well plates. Lentivirus were packaged by co-transfection of the transfer plasmid encoding the protein of interest with psPAX2 (Addgene #12260) and pVSVG (Addgene # 35616) in 293FT cells (ThermoFisher), and the media containing virus was collected ∼48 h later, centrifuged to remove debris, and used directly for infection. Retrovirus were packaged similarly but by transfection of the transfer plasmid into 293 Phoenix Ampho cells. All cell lines were routinely checked to be mycoplasma-free by DAPI staining.

U2OS REEP4 knockout cells were generated using CRISPR and purified AsCas12a/AsCpf1. AsCas12a/AsCpf1 protein was purified as described^47^ from an expression plasmid generated by deleting the MBP sequence from plasmid pDEST-hisMBP-AsCpf1-EC (Addgene plasmid # 79007) transformed into Rosetta(DE3)pLysS Competent Cells (Novagen). 80 pmol of Alt-R CRISPR-Cas12a crRNA targeting sequence GGATGCTGTGTCCAGCTTATGCTT (IDT) was incubated with 63 pmol of AsCas12a/AsCpf1 protein and electroporated into 2×10^5^ U2OS cells along with 39 pmol of Alt-R Cpf1 Electroporation Enhancer (IDT). Isolated clones carrying deletions in all REEP4 gene copies were identified by PCR amplification of the targeted region with primers 5’-GTACAAGCCAAGCGAGAAAAAC and 5’-CTATTAAACACACCCAAACCCC and Illumina MiSeq, and further confirmed by immunoblotting with αREEP4 antibodies.

For transient plasmid transfections, cells were plated into 6-well plates and transfected at ∼75% confluency with1 μg of DNA using lipofectamine 3000 (ThermoFisher). For imaging, cells were trypsinized and replated onto No 1.5 cover glass (Warner Instruments) the following day and analyzed ∼12-16 h later by indirect immunofluorescence. For starvation experiments, cells were triple washed and incubated with EBSS (Millipore Sigma #E3024) for thirty min before fixation. For drug treatments, cells were incubated with nocodazole (50 μM, 1 h; Millipore #487928) or Brefeldin A (20 μM, 1 h; Millipore #203729) before fixation and immunofluorescence analysis. For cycloheximide chase experiments, cells were split equivalently into two 2 cm^2^ wells 12-16 h post-transfection and analyzed a day later.

For protein purification, 50 ml of Expi293 suspension cells at ∼2.2 x 10^6^ cells/ml were transiently transfected with 50 μg of total plasmid DNA encoding human ATL1 tagged with a C-terminal FLAG tag, and/or human REEP1 tagged with a C-terminal SBP tag, using 150 μl of 1mg/ml PEI Max (Polysciences #24765). Expression was boosted with the addition of 10 mM sodium butyrate (Millipore) ∼16 h post transfection, and cells were collected 48 h later for purification.

### Indirect immunofluorescence and confocal fluorescence microscopy

For indirect immunofluorescence, U2OS cells were grown on acid washed No 1.5 glass coverslips (Warner Instruments), fixed in PBS/4% paraformaldehyde (Electron Microscopy Services), and permeabilized with PBS/0.1% TritonX-100 for 5 min. Samples were blocked in PBS/1% FBS/0.01% Triton X-100 for ∼1 h, and then sequentially probed with primary and secondary antibodies conjugated to Alexafluor dyes (405, 488, 568, or 647, from ThermoFisher) with 3x PBS washes after each primary and secondary antibody incubation. Samples were mounted onto glass slides with fluoromount-G (Southern Biotech). To visualize F-actin, cells were incubated with phalloidin-405 after the secondary antibody staining, washed, and mounted as above. For neutral lipid staining, cells were grown in glass-bottom Mattek dishes, and after secondary antibody staining, lipidtox Deep Red was added at 1:1000 dilution in PBS and imaged directly.

Confocal fluorescence microscopy was performed on a Nikon Ti inverted microscope equipped with a Hamamatsu ORCA-Fusion BT sCMOS camera, Nikon LUN-F XL solid state lasers (405, 488, 561, and 640 wavelengths), and a 100X Plan Apo 1.4 N/A oil objective. Images were acquired using either Metamorph or NIS elements software, and FIJI/ImageJ was used for all image analysis and processing. For display, fluorescence intensities of raw 16-bit depth images were linearly scaled across the entire image and then converted to 8 bit. Images showing cropped enlargements were similarly processed. Multicolored overlays were pseudocolored.

To correlate REEP1-HA expression levels with different localization phenotypes, images were taken with a 63X 1.4 N/A oil objective using identical exposure settings for the anti-HA fluorescence channel, and the average fluorescence intensity, subtracted for background, of each cell was compared to its localization. For Pearson’s coefficient calculations, images were taken using a 100X Plan Apo 1.4 N/A oil objective. A ∼9×9 μm area at the cell periphery, where the ER is a clear monolayer, was selected and analyzed for colocalization using the JaCoP/Biop JaCoP plugin. All quantifications were performed on raw 16-bit images, and calculations, statistical analysis and graphs were analyzed and plotted using Microsoft Excel and Graphpad Prism.

### Thin-section electron microscopy to visualize APEX-DAB labeled REEP1 vesicles

U2OS cells stably expressing REEP1-APEX were fixed for 1 h in PBS containing 2% paraformaldehyde/1% glutaraldehyde, quenched with 20 mM glycine for 20 min, and then washed 5x in 100 mM sodium cacodylate buffer. Samples were then incubated with 0.5 mg/ml diaminobenzidine (Millipore #D8001)/0.1 M HCl and 10 mM H_2_O_2_ (Millipore #H1009) for 1 h on ice, and washed 5x in sodium cacodylate buffer. Samples were then fixed and stained with 1% OsO_4_ reduced with 1.5% ferrocyanide, dehydrated sequentially in 20%, 50%, 70%, 90%, and 100% (vol/vol) ethanol on ice, and infiltrated and embedded in Epon resin. Ultrathin sections (∼70 nm) were cut on a Reichert Ultracut-S microtome and mounted onto copper grids. Images were acquired on a JEOL 1200X transmission electron microscope equipped with an AMT 2k CCD camera.

### Membrane fractionation and cycloheximide chase experiments

For fractionation, ∼5×10^6^ U2OS cells stably expressing REEP1-mEm or mEm-REEP5 were washed twice in PBS containing 1 mM phenylmethylsulfonyl fluoride (PMSF), twice with hypotonic buffer (20 mM HEPES pH 7.4, 10 mM KCl, 1 mM EGTA, 1 mM PMSF), and then collected in 1 ml of hypotonic buffer by scraping. Cells were lysed in a glass douncer, and lysates were centrifuged at 1,000 xg for 2 min to remove unbroken cells and nuclei. KCl was added to 150 mM, and lysates were centrifuged at 10,000 xg for 5 min to pellet mitochondria. 50 μl of this clarified lysate sample was collected as ‘Input’ and the rest divided into two tubes and fractionated by ultracentrifugation at 100,000 xg for 1 h at 4 °C. The supernatant containing soluble proteins was collected (‘S/N’), and the pellets were washed in buffer containing 0.5 M KCl to remove peripherally bound proteins to the membrane and centrifuged again at 100,000 xg for 30 min. One tube pellet was directly denatured with 2x Laemmli buffer for analysis (‘pellet’), while the other was solubilized in buffer containing 1% dodecyl maltoside for 1 h before a final centrifugation step at 100,000 xg for 1 h. The resultant supernatant (‘pellet +DDM’) was collected and denatured with Laemmli buffer for immunoblot analysis.

For cycloheximide chase analysis of REEP1-HA constructs, transiently transfected U2OS cells were treated with 50 μM cycloheximide for 6 h before lysing cells in modified RIPA buffer (50 mM HEPES pH 7.4, 150 mM NaCl, 1 mM MgCl_2_, 1% TritonX-100, 0.1% deoxycholate, 0.1% SDS, with protease inhibitors). Lysates were clarified by centrifugation at 15,000 rpm for 10 min, denatured in Laemmli buffer, and analyzed by immunoblotting with anti-HA and anti-tubulin antibodies.

Immunoblotting was performed using standard procedures: Samples were resolved on a 4-20% gradient TGX-PAGE gel (BioRad) and transferred onto nitrocellulose. Membranes were blocked in 1-2% milk/PBS and incubated with primary antibodies and HRP-conjugated secondary antibodies with triple wash steps after each incubation. Immunoblots were developed using western lightning ECL (Perkin Elmer) and imaged on an Amersham 800 Imager (Cytiva).

### Protein purification of REEP1-SBP/ATL1-FLAG complexes

Transfected Expi293 cell pellets were solubilized in 3% digitonin in 50 mM HEPES pH 7.5, 800 mM NaCl, supplemented with DNase I and protease inhibitors, for 1.5 h. The lysate was centrifuged at 4 °C for 15 min in a TLA55 rotor (Beckman) at 50,000 rpm to remove insoluble material. For purification of REEP1-SBP alone, the clarified supernatant was incubated with streptavidin agarose resin (GoldBio), eluted in buffer containing 5 mM biotin, and further purified by size exclusion chromatography on a Superose 6 column (Cytiva) in 20 mM HEPES pH 7.5, 150 mM NaCl, and 0.06% digitonin. For ATL1-FLAG and REEP1-SBP copurification, a similar protocol was followed except the clarified supernatant was incubated with anti-FLAG resin (Millipore) and eluted with 0.2 mg/ml 3xFLAG peptide (Millipore). All samples were denatured in Laemmli buffer and analyzed by SDS-PAGE (4-20% TGX, Biorad) and Coomassie blue staining.

## Supplementary Figure Legends

Supplementary Figure 1. REEP1 proteins localize to punctae independent of tags, expression levels, or cell line.

**a,** U2OS cells stably expressing the ER marker RFP-Sec61β were transfected with HA-tagged REEP1, REEP2, REEP3, or REEP4 and analyzed by indirect immunofluorescence with anti-HA (αHA) antibodies and confocal fluorescence microscopy. Scale bars, 2 µm.

**b.** As in a, but of a cell expressing high levels of REEP1-HA that has a bundled tubular localization. Left column shows a maximal projection of a z-series through the entire cell volume. Right column shows a magnified, single focal plane view of the boxed region. Scale bar, 10 µm.

**c**, Quantification of REEP1-HA localization compared to relative expression levels of transiently transfected cells, as analyzed in a. The average fluorescence intensity of αHA across the cell was compared to relative localization patterns of punctae only, punctae and bundled tubules, or predominantly bundled tubules.

**d**, As in a, but in a U2OS cell transfected with untagged REEP1 and immunostained with antibodies against REEP1 (αREEP1) and the general ER membrane marker calnexin (αCNX). Right panels show magnifications of the boxed region. Scale bar, 2 µm.

**e**, REEP1 punctae localization was measured across expression levels. Pearson’s correlation coefficients (r) were quantified comparing αREEP1 and αCNX colocalization of cells as in d, and r values were plotted against the mean intensities of the anti-REEP1 signal across the whole cell. n, 151 cells.

**f**, As in d, but in a cell transfected with an untagged REEP2 construct and immunostained with αREEP2 antibodies. Scale bar, 2 µm.

**g**, As in d, but transfected with an untagged REEP4 construct and immunostained with αREEP4 antibodies.

**h,** Hela cells stably expressing human REEP1, REEP4, or REEP5 fused to mEmerald (REEP1-mEm, REEP4-mEm, REEP5-mEm) were immunostained with antibodies against the ER luminal marker αKDEL. Scale bar, 2 µm.

**i**, Endogenous REEP2 localization was determined by immunostaining with αREEP2 antibodies and compared to calnexin in neuronal SKN-SH cells. Scale bar, 2 µm.

**j**, Endogenous REEP4 localization in U2OS cells was determined by immunostaining with αREEP4 and compared to αCNX. Middle row shows magnified views of the boxed region. REEP4 antibodies are specific to REEP4, as shown by immunostaining of U2OS CRISPR-knockout cells lacking REEP4 (REEP4 KO, bottom row) and imaged using identical exposure settings as the parental line. Scale bars, whole cells, 10 µm; magnification, 2 µm.

Supplementary Figure 2. Mutations in the C-terminal APH relocalize REEP1 to the bulk ER.

**a**, Constructs encoding HA-tagged REEP1 mutants with different C-terminal deletions (Δ82-95, Δ100-139, or Δ156-201) were transfected into U2OS cells stably expressing RFP-Sec61β, and their localization was determined by immunostaining with αHA antibodies and confocal microscopy. Pearson’s correlation coefficients (r) between αHA and RFP-Sec61β localizations are shown in the inset. Scale bar, 2 µm.

**b**, As in a, but with REEP1 containing single lysine mutations (I103K, L107K, A110K, F121K, and A132K) introduced into the APH-C.

**c**, As in a, but with REEP1 containing point mutations (L96P, S97P, A110E, and S114N) that have been linked with disease and relocalize REEP1 to the bulk ER.

**d**, As in a, but with HSP-linked point mutations that do not affect REEP1 localization to vesicles.

Supplementary Figure 3. The TM domain of REEP1 is important for vesicle localization.

**a,** Schematic of the TM domains of REEP1. Primary sequence conservation was analyzed using ConSurf (http://consurf.tau.ac.il), and disease mutations in HSP are indicated. TM segments are marked in blue, above.

**b**, U2OS cells stably expressing RFP-Sec61β were transfected with a HA-tagged REEP1 construct carrying the P19R disease mutation, and localization was determined by αHA immunostaining and confocal microscopy. Lipid droplets (LD) were visualized with Lipidtox. Arrowheads point to examples of colocalized αHA and LD signals. Top row shows a cell with numerous, small REEP1(P19R)-HA punctae; the bottom row shows a cell with REEP1(P19R)-HA at the rim of a larger LD. Scale bar, 2 µm.

**c**, As in b, but with a REEP1-HA construct carrying the P19L disease mutation.

**d**, REEP1-HA constructs carrying the A20E, S23F, W42R, T55K, D56N, or no (WT) disease mutation were transfected into U2OS cells and analyzed as in b. Shown are the merged images of αHA and lipidtox signals.

**e**, The relative stabilities of REEP1 TM mutants were analyzed by cycloheximide chase experiments. Cells transfected with equivalent DNA amounts of wildtype or mutant REEP1-HA were collected at 0 or 6h after cycloheximide (CHX) treatment, and lysates were analyzed by immunoblotting with αHA antibodies. Anti-α tubulin serves as a loading control. The middle blot shows the HA blot linearly adjusted for gain (oversat.) to visualize the dimmer REEP1-HA mutant bands.

**f-l**, U2OS cells stably expressing REEP1-mEm were transfected with REEP1-HA carrying the indicated disease mutation, and localization was analyzed by anti-HA immunostaining and confocal fluorescence microscopy.

**m**, As in f-l, but with wildtype (WT) REEP1-HA.

**n**, Pearson’s correlation coefficients were measured between the αHA and mEm signals of samples as analyzed in f-m. Shown are means and standard deviations. n, 65-100 cells/sample.

**o**, Alphafold-predicted model of the REEP1 dimer interface. Residues whose mutation causes destabilization and relocalization are highlighted in yellow.

Supplementary Figure 4. REEP1 vesicles are distinct from other organelles and insensitive to inhibition of ER to Golgi transport.

**a**, A U2OS cell stably expressing REEP1-mEm and RFP-Sec61β was stained with phalloidin to visualize F-actin and imaged by confocal microscopy. Bottom row shows magnifications of the boxed region. Scale bar for whole cell, 10 μm; magnification, 2 μm.

**b**, As in a, but with a cell immunostained with anti-α tubulin antibodies to visualize microtubules.

**c**, As in b, but after 30 min treatment with the microtubule polymerization inhibitor nocodazole.

**d**, A U2OS cell stably expressing REEP1-mEm was immunostained with Tom20 antibodies to visualize mitochondria. Scale bar, 2 μm.

**e**, As in d, but with immunostaining with antibodies against giantin to visualize the Golgi. Bottom row shows magnifications of the boxed region. Scale bars for whole cell, 10 μm; magnification, 2 μm.

**f**, As in e, but in a cell also stably expressing RFP-Sec61β and treated with Brefeldin A to inhibit trafficking between ER and Golgi. Note that this inhibitor also disperses the Golgi.

**g**, As in f, but with a U2OS cell stably expressing REEP1-mEm and RFP-Sec61β grown at 16 °C for 16 h to block vesicle trafficking through the secretory pathway. Scale bar, 2 μm.

**h**, A U2OS cell stably expressing REEP1-mEm and transfected with mCherry-DFCP1 (mCh-DFCP1), starved in EBSS for 30 min, and immunostained with anti-LC3 (αLC3) and anti-LAMP1 (αLAMP1) antibodies. DFCP1 marks sites for autophagosome formation on the ER, LC3 marks phagophore membranes, and LAMP1 is a lysosome marker. The merged image shows the overlay of REEP1-mEm, mCh-DFCP1, and αLC3. Right panels show magnifications of the boxed regions. Scale bar for whole cell, 10 μm; magnifications, 1 μm.

Supplementary Figure 5. Interaction between REEP1-SBP and ATL1-FLAG.

Expi293 cells were transfected with REEP1-SBP alone or with ATL1-FLAG, and digitonin-solubilized cell lysates were subjected to immunoprecipitation with beads containing anti-FLAG antibodies. The beads were washed twice (wash 1, 2) and bound material eluted with FLAG peptide in four fractions. All samples were analyzed by SDS-PAGE and Coomassie blue staining. Note that REEP1-SBP co-elutes with ATL1-FLAG.

## Notes

### Competing Interest Statement

The authors have declared no competing interest.

