## Supplementary Figs. 1-5 for "The membrane curvature inducing REEP1 proteins generate a novel ER-derived vesicular compartment"

#### Supplementary Figure 1

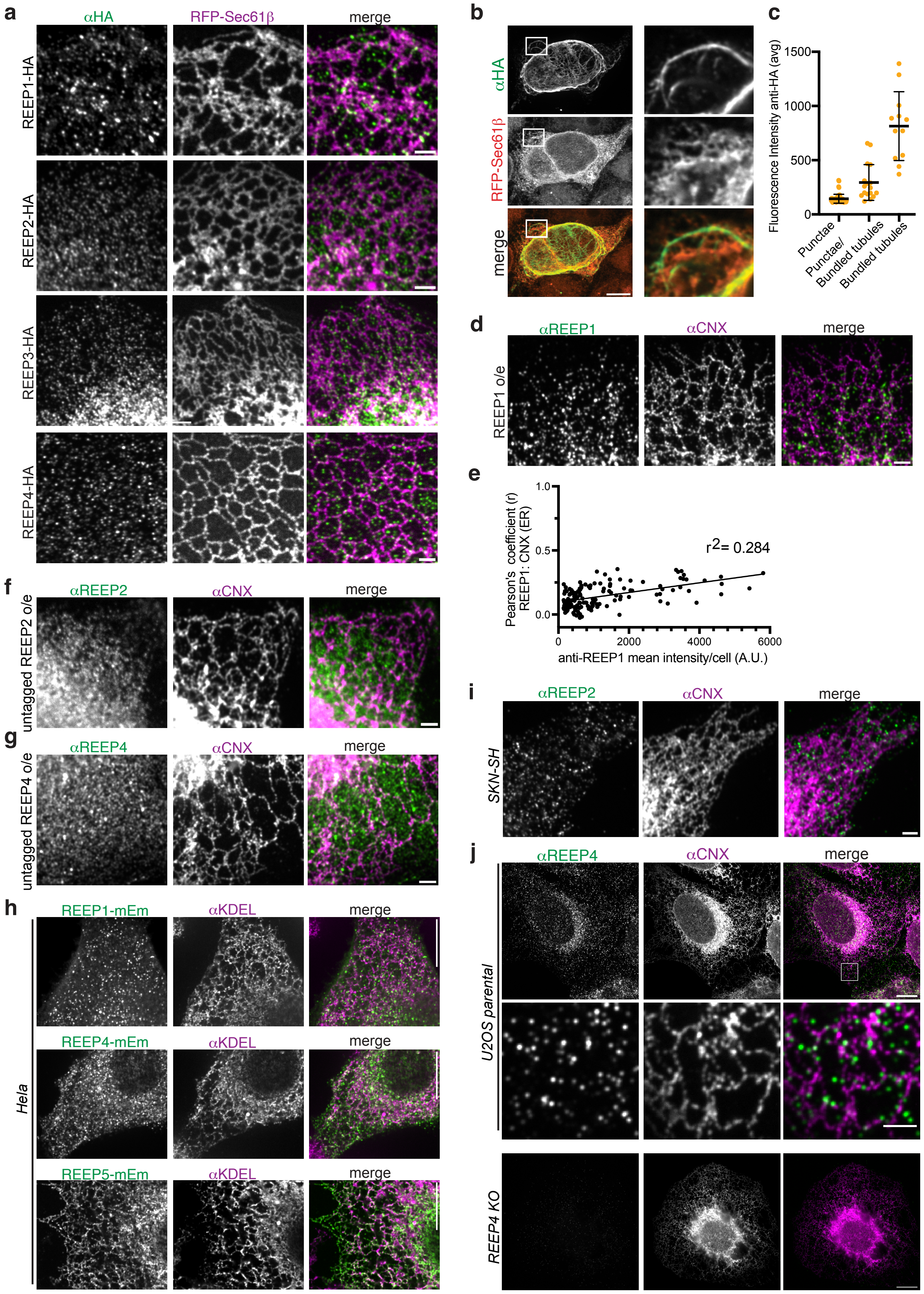

**Supplementary Figure 2**

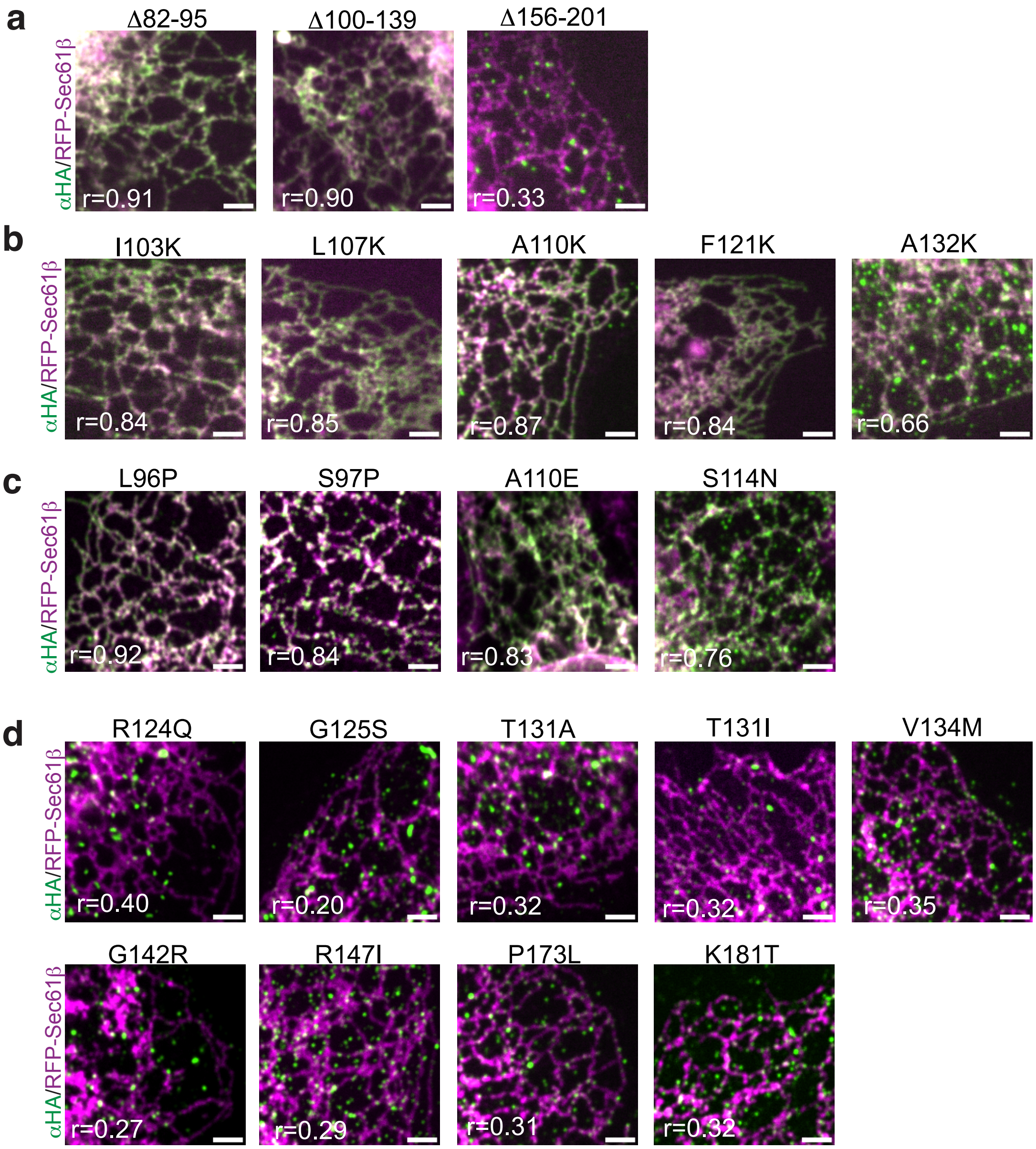

#### Supplementary Figure 3

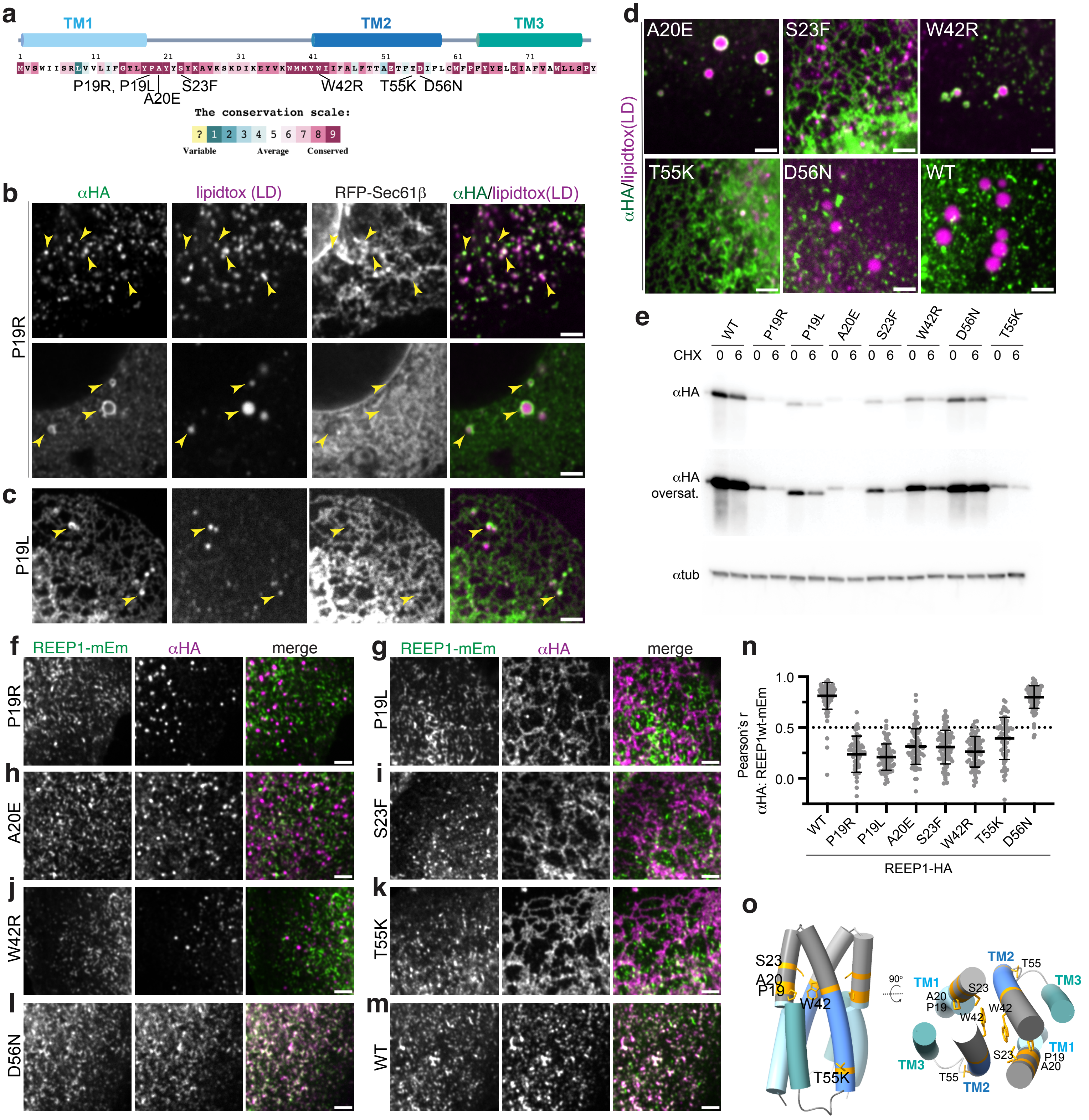

### Supplementary Figure 4

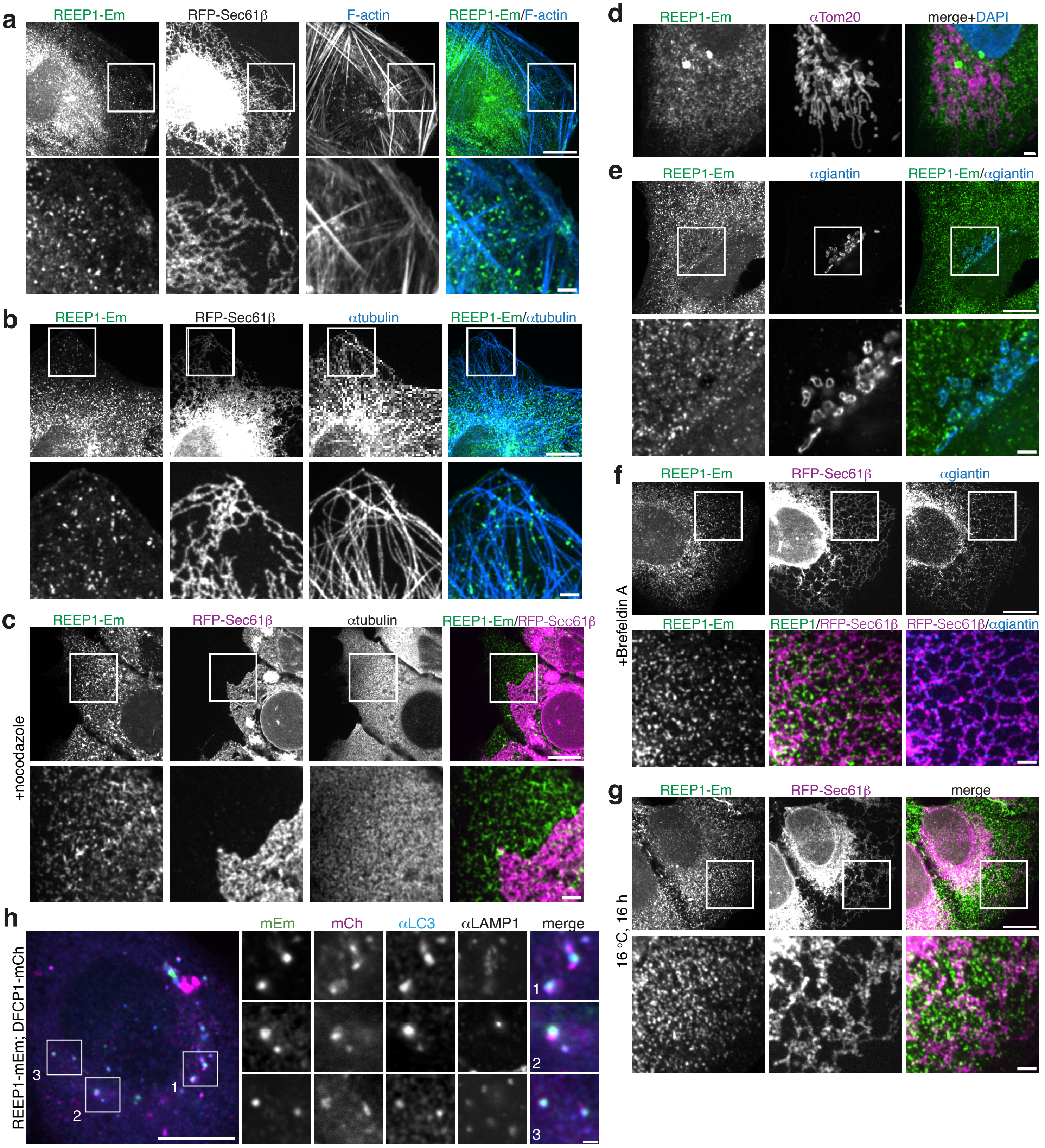

#### Supplementary Figure 5

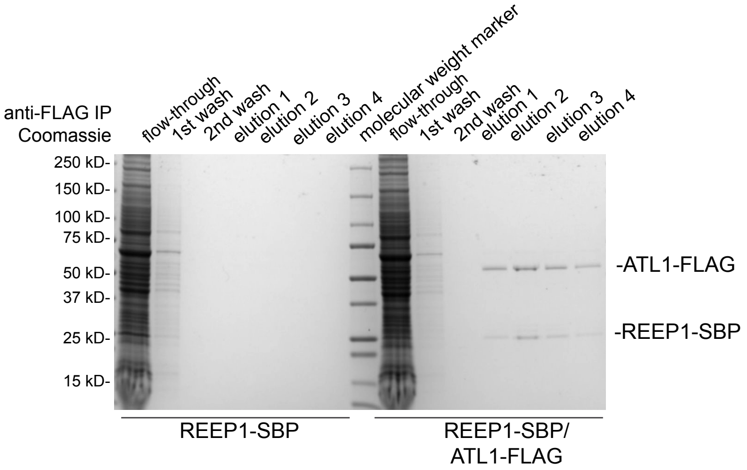
